# Reciprocal constraint couples architectural protein abundance and pericentromeric satellite expansion

**DOI:** 10.64898/2026.05.15.725447

**Authors:** Damian Dudka, Keagan Beeravolu, Michael A. Lampson

## Abstract

Long-read sequencing has enabled precise measurements of highly repetitive centromeric satellites and their rapid divergence between species^1–7^. Large satellite arrays emerge from libraries of shorter arrays via stochastic expansions^8,9^, but understanding the selective pressures constraining such expansions remains a major challenge. Here, using the mouse “major” satellite as a model, we reveal reciprocal functional constraints between increasing satellite copy number and abundance of a conserved architectural protein in female meiosis. We show that HMGA2 (high mobility group AT-hook 2) is enriched at major satellite, and its expression correlates with major satellite copy number: both are high in *Mus musculus* compared to the closely related *Mus spretus*. To test functional constraints, we modulated HMGA2 abundance by depletion or overexpression and used a *musculus*/*spretus* hybrid to generate oocytes with intermediate HMGA2 expression and major satellite copy number. We find that HMGA2 depletion disrupts major satellite packaging in major satellite-rich *musculus* but not hybrid oocytes, indicating that increasing copy number requires high HMGA2 expression. Conversely, HMGA2 overexpression disrupts chromosome segregation in major satellite-poor *spretus* but not hybrid oocytes, indicating that high HMGA2 expression requires expanded major satellite arrays. Based on these results, we propose a co-evolution model in which satellite expansion is constrained by architectural protein abundance, whereas protein abundance is constrained reciprocally by satellite array size.

## Results

Centromeres are specialized chromatin regions mediating the conserved process of chromosome segregation. Paradoxically, centromeres are typically built on highly repetitive and rapidly evolving DNA satellites that differ in sequence and array size between closely related species^10–12^. Although centromeres and pericentromeres have key epigenetic components (the histone H3 variant CENP-A and H3K9me3 heterochromatin, respectively)^13–15^, accumulating evidence suggests that divergence of the underlying satellite DNA has functional consequences. Centromeric and pericentromeric chromatin organization and packaging differ across divergent satellites^16–18^, which can lead to reproductive isolation^17,19–21^. Moreover, differences in centromeric satellite array sizes are associated with Down syndrome^22^ and are prevalent in cancer^23–25^.

Satellites originate from preexisting tandemly repeated sequences such as rDNA, simple repeats, and transposon-derived repeats^26,27^. Under the library hypothesis, closely related species inherit an ancestral “library” of small arrays from which larger arrays emerge via independent expansions^8^. Consistent with this hypothesis, clade-specific satellite libraries were found in plants^28^, insects^9,29,30^, fish^31,32^, and mammals^12,33^. Furthermore, by resolving large arrays and tracing array ancestry, long-read sequencing supported the emergence of large arrays from smaller ones^2,28^. Several centromeric satellite expansion mechanisms have been proposed^34^ ^35^, but whether these expansions are stochastic or selection-driven has been an ongoing debate^36–38^.

Early simulations suggested that unequal crossovers without selection are sufficient to explain satellite expansion and divergence^39,40^. While centromeric satellites can expand in cancer cell lines passaged without selection^35^, genomic and population genetics analyses indicate that stochastic expansions do not fully account for the evolutionary dynamics of satellites across species^41–44^. Structural properties of centromeric satellite DNA are consistent with selective constraints. For instance, biased nucleotide composition of fly satellites might enhance nucleosome positioning and recruit sequence-dependent proteins^42^. In mammals, centromeric satellites contain dyad symmetries, form non-B DNA, and are enriched for a motif that recruits a centromere protein, CENP-B^45,46^. Once expanded, some centromeric satellites show a selective advantage by propagating selfishly through female meiosis^47–49^, a phenomenon known as centromere drive^50,51^, despite imposing a fitness cost^52,53^. These findings support the role of selection in satellite evolution but do not explain the expansion of divergent satellites in related species.

Two closely related mouse species (*Mus musculus* and *Mus spretus*) are a good model for lineage-specific satellite expansion because they carry two divergent satellites at centromeric regions that expanded either in *musculus* (major satellite) or *spretus* (minor satellite). Major satellite is packaged during female meiosis by condensin II^17^ and the DNA shape-recognizing architectural protein HMGA1^18^. Major satellite abundance correlates with condensin II across mouse species, suggesting a co-evolution between major satellite and condensin^17^. However, condensin is largely depleted from major satellite^17,48^, whereas HMGA1 is enriched^18^. Furthermore, the depletion phenotypes suggest that HMGA1 is the dominant packaging protein for large major satellite arrays. We recently showed that major and minor satellites differ in DNA shape (minor groove width) and hypothesized that lineage-specific expansion could be due to differences in DNA shape-recognizing proteins between closely related species^18^ (**Fig. 1A**). Since HMGA1 sequence and abundance do not differ between *musculus* and *spretus*^18^, we searched for other candidates. In this paper, we show that another conserved DNA shape-recognizing architectural protein, HMGA2, is more highly expressed in *musculus* vs *spretus* and required to package the increased major satellite copy number in *musculus*. We also find that high HMGA2 expression interferes with chromosome segregation in major satellite-poor *spretus*, revealing reciprocal functional constraint between HMGA2 abundance and major satellite copy number.

**Fig. 1.**
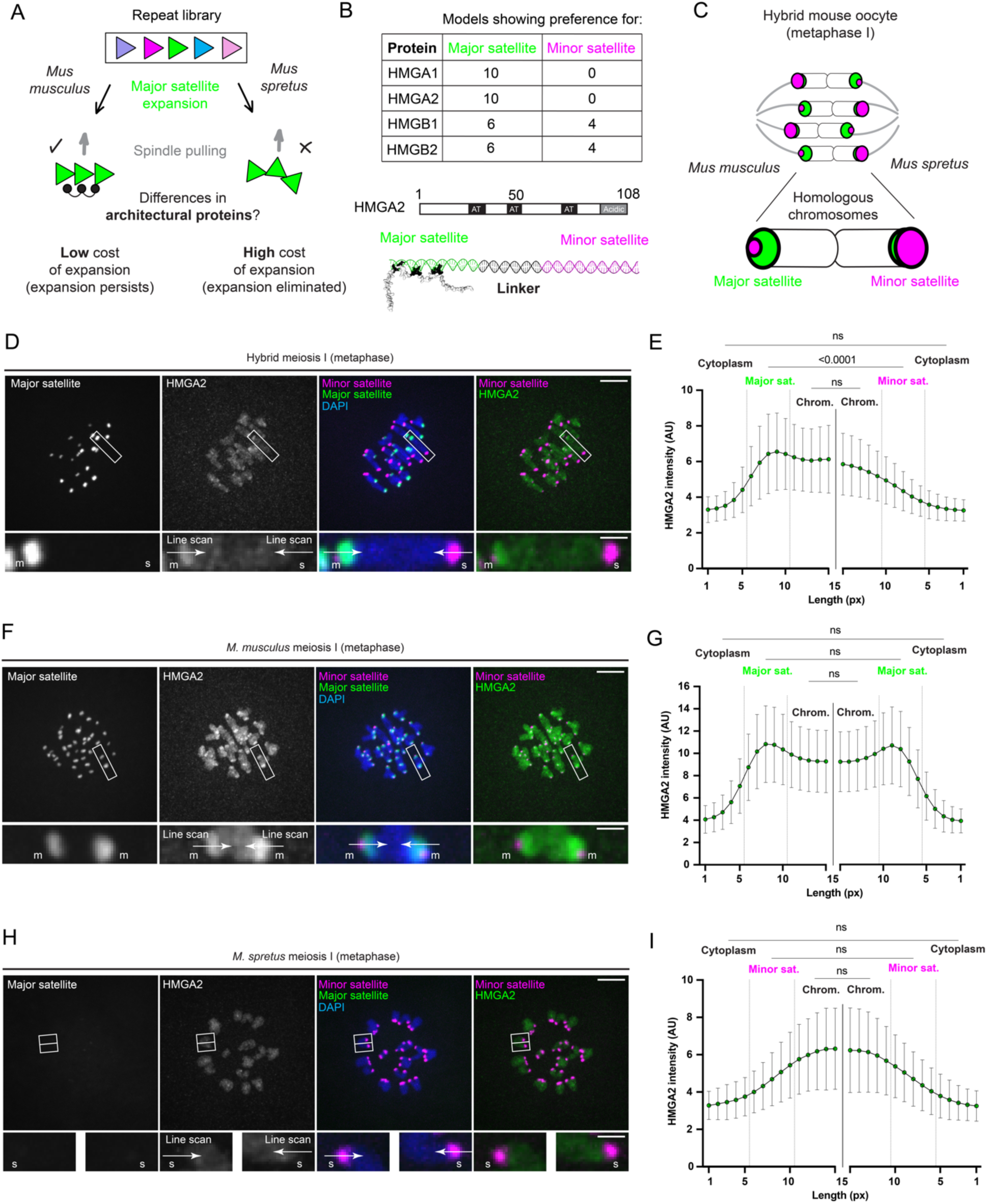
HMGA2 preferentially binds to major satellite. **A**) We hypothesize that rapid expansion of a centromeric satellite from a library (colorful triangles) is tolerated only in the presence of a DNA shape-recognizing architectural protein (black circles) that packages the expanded array to withstand microtubule pulling forces (grey arrows). **B**) The table (top) summarizes AlphaFold3 modeling results, and the structural model (bottom) shows HMGA2 binding preference for major (green) over minor (magenta) satellite fragments, separated by a poly-guanine linker (black). AT-hook motifs (colored black in the structure prediction) and the acidic tail are indicated in the HMGA2 amino acid sequence. **C**) Schematic of a *musculus*/*spretus* hybrid oocyte at metaphase I, showing microtubules (grey), major satellite (green), and minor satellite (magenta). **D-I**) Images show hybrid (D), *musculus* (F), or *spretus* (H) oocytes in meiosis I, expressing TALE-mClover targeting major satellite and dCas9-mCherry with gRNA targeting minor satellite, fixed and stained with anti-HMGA2 antibody. Arrows show line scans along homologous chromosomes. Scale bars: 10 μm or 2 μm (inset); m: *musculus*, s: *spretus*. Plots show quantification of HMGA2 intensity in hybrid (E), *musculus* (G), and *spretus* (I) oocytes, averaged over multiple line scans. Bars show standard deviations; n=230-300 line scans from 23-30 oocytes from 3 independent experiments; Kruskal-Wallis test followed by Dunn’s multiple comparisons test, ns: not significant.

### HMGA but not HMGB proteins preferentially bind to major satellite

To identify architectural proteins that recognize the DNA shape of major satellite, we designed a simple *in silico* DNA shape-recognition assay using AlphaFold3 (**Fig. 1B**). We combined major and minor satellite repeat fragments separated by a poly-guanine linker DNA and modeled the binding of architectural proteins from the minor groove-binding HMG superfamily: HMGA1, HMGA2, HMGB1, and HMGB2. HMGA proteins contain three AT-hook motifs that recognize narrow DNA minor grooves of AT-stretches^54^ (contiguous A and T bases) and crosslink multiple DNA molecules^55^. HMGB proteins bind to the DNA minor groove via two HMG boxes and bend DNA^56,57^. As expected, HMGA1 showed a clear preference for binding to major over minor satellite across all prediction models (**Fig. 1B**). HMGA2 also preferred binding to major satellite (**Fig. 1B**), but HMGB1/2 did not have any preference (**Fig. S1A**). To test the AlphaFold3 predictions, we stained endogenous HMGA2 in *musculus*/*spretus* hybrid oocytes in meiosis I, when homologous chromosomes with divergent satellites are paired (**Fig. 1C**). As predicted, HMGA2 was enriched at major satellite compared to minor satellite (**Fig. 1D and E**). Consistent with this result, we observed HMGA2 enrichment at major satellite in the parental *musculus* oocytes (**Fig. 1F and G**), but not at minor satellite in *spretus* (**Fig. 1H and I**). Since endogenous HMGB1 and HMGB2 cannot be detected during chromosome segregation using immunostaining due to transient chromatin association^58^, we expressed GFP-tagged constructs and imaged cells live. Consistent with the AlphaFold3 predictions, we did not observe HMGB1 or HMGB2 enrichment at either satellite in hybrid oocytes (**Fig. S1B and C**). These results show that major satellite DNA shape is recognized by architectural proteins containing AT-hook motifs but not HMG boxes.

### HMGA2 expression correlates with the major satellite copy number

Our satellite expansion hypothesis predicts differences in architectural protein sequence or abundance correlating with divergent satellite expansions in closely related species (**Fig. 1A**). The HMGA2 sequence is the same between *musculus* and *spretus*, but immunofluorescence suggested differences in HMGA2 chromatin abundance across species (**Fig. 1D-I**). We found that HMGA2 intensity was twice as high at major satellite in *musculus* oocytes as in hybrids (**Fig. 2A**), and also significantly reduced at chromosome arms in *musculus* compared to *spretus* or hybrid oocytes (**Fig. 2B**). We further tested HMGA1 and HMGA2 expression levels by RT-qPCR and found that HMGA2 transcript levels in *musculus* oocytes were 2-fold higher than in hybrid oocytes and 4-fold higher than in *spretus* (**Fig. 2C**). In contrast, HMGA1 transcript levels were similar between species (**Fig. 2D**), consistent with our previous immunofluorescence data^18^. Together, these results show that expression differences lead to differential HMGA2 protein abundance at chromatin. We conclude that, unlike HMGA1, HMGA2 expression correlates with major satellite copy number, consistent with our satellite expansion hypothesis (**Fig. 1A**).

**Fig. 2.**
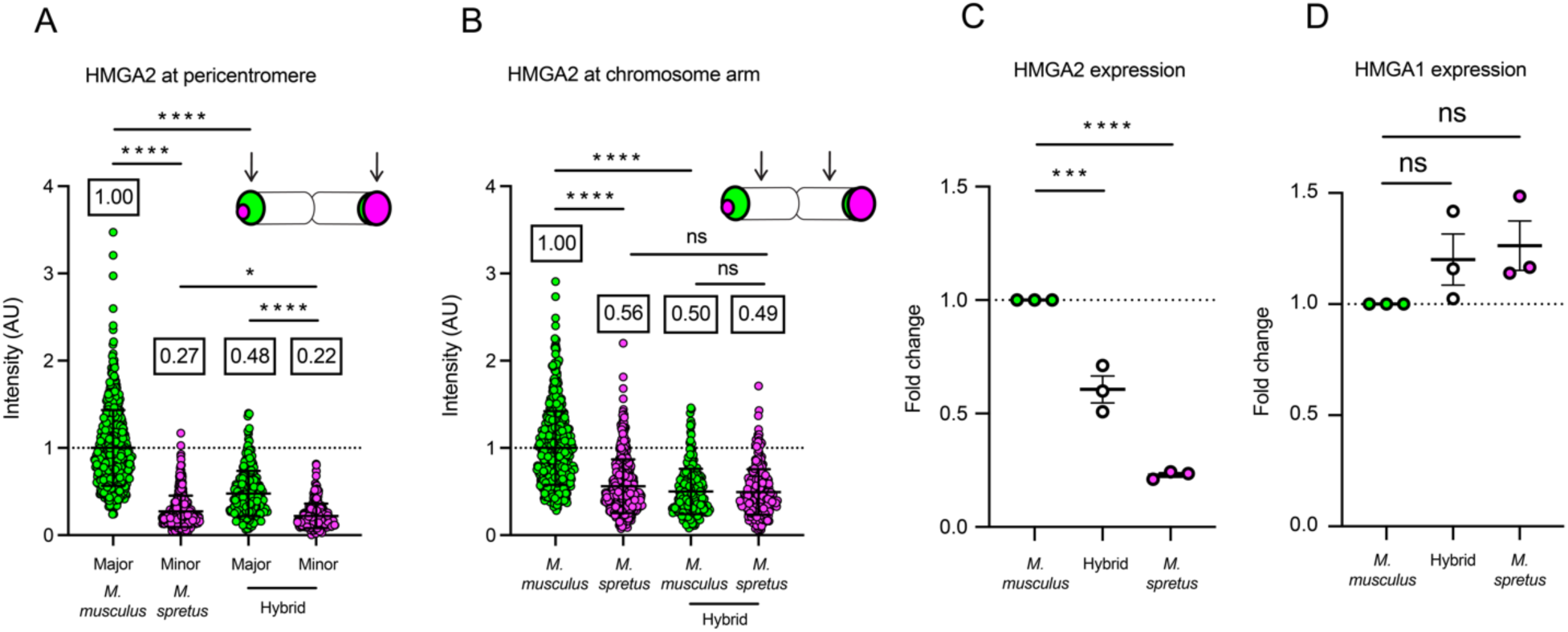
HMGA2 expression correlates with major satellite copy number. **A** and **B**) Relative HMGA2 protein levels at *musculus*, hybrid, and *spretus* pericentromeres (A) or chromosome arms (B) based on data represented in Fig. 1D, F and H. Each dot represents a single pericentromere; bars show mean and standard deviation; n=230-300 measurements from 23-30 oocytes from 3 independent experiments; Kruskal-Wallis test followed by Dunn’s multiple comparisons test, *p<0.05, ****p<0.0001, ns: not significant. **C** and **D**) Relative HMGA2 (C) and HMGA1 (D) mRNA levels using RT-qPCR in *musculus*, hybrid, and *spretus* GV oocytes. Each dot represents mRNA pooled from 30 oocytes; bars show mean and standard error of the mean; N=3 experiments, n=30 oocytes per experiment; one-way ANOVA test followed by Dunnett’s multiple comparisons test, ***p<0.001, ****p<0.0001, ns: not significant.

### HMGA2 depletion disrupts major satellite packaging in *musculus* but not hybrid oocytes

HMGA1 and HMGA2 diverged in the last common ancestor of vertebrates and show very little global sequence similarity^59^. Nevertheless, their AT-hooks are nearly identical^60^, suggesting that both could contribute to major satellite packaging. We previously showed that acute antibody-mediated degradation of HMGA1 disrupts major satellite packaging in both *musculus* and hybrid oocytes^18^. Using the same approach (**Fig. 3A-C** and **Fig. S2**), we find that HMGA2 depletion has little impact on minor satellite packaging in hybrid or *spretus* oocytes, consistent with the lack of enrichment at these sequences (**Fig. 1**). HMGA2 depletion also has no meaningful effect on major satellite packaging in hybrid oocytes, indicating that HMGA1 is sufficient in these cells. The ∼2-fold difference in major satellite copy number between *musculus* and hybrids allowed us to test whether HMGA2 becomes required as major satellite copy number increases. We find that HMGA2 depletion in *musculus* oocytes leads to massive major satellite stretching, similar to the effects of HMGA1 depletion but without loss of HMGA1 (**Fig. 3D and E**). Comparing the length of major satellite in HMGA2-depleted *musculus* and hybrid oocytes revealed a nearly 4-fold difference (**Fig. 3C**). These results suggest that HMGA1 is sufficient to package half of the major satellite currently found in *musculus*, but two-fold satellite expansion in the *musculus* lineage (7.5-10% of the entire genome^5,6,18^) required high HMGA2 abundance to withstand spindle forces in female meiosis.

**Fig. 3.**
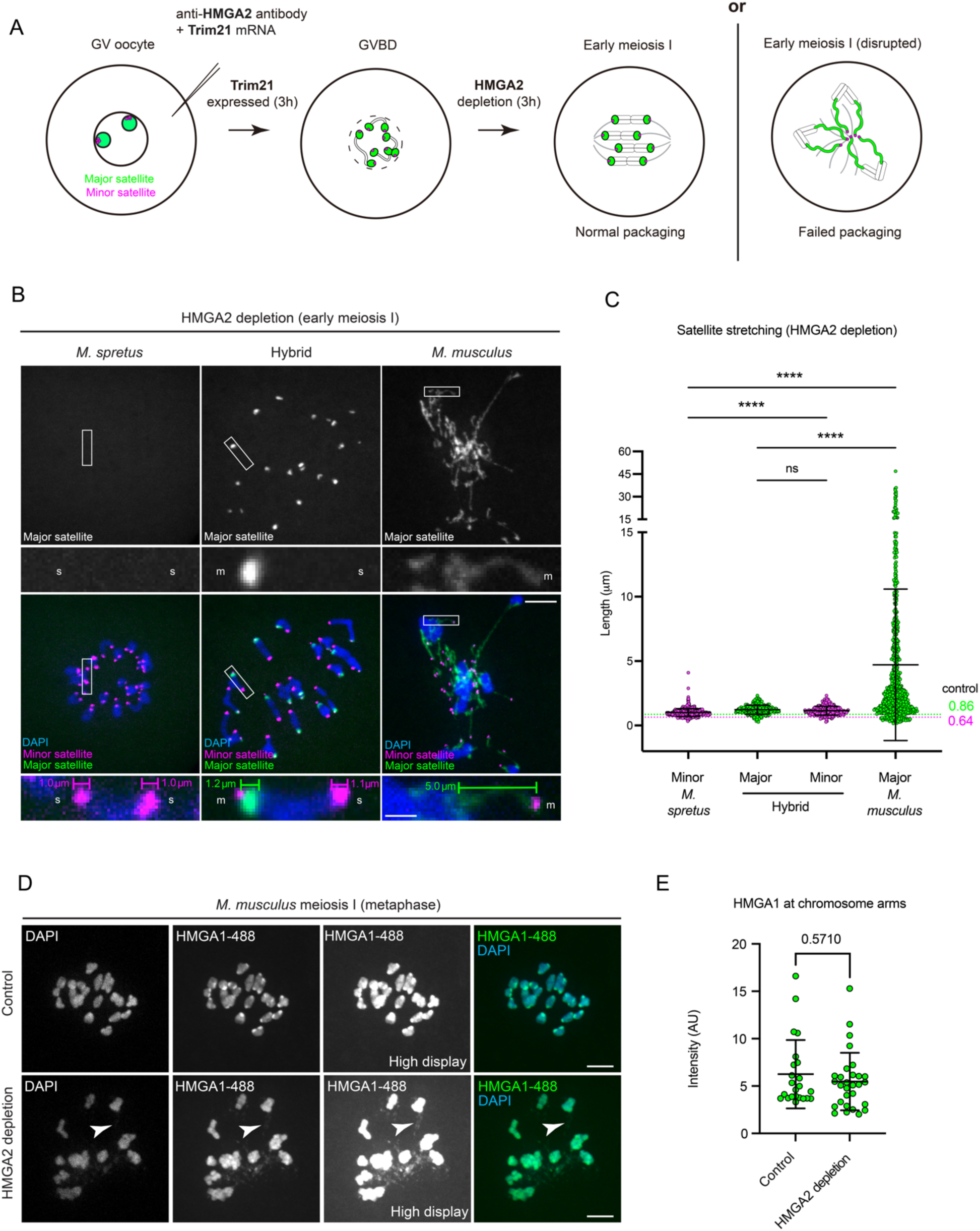
HMGA2 depletion disrupts major satellite packaging in *musculus* but not hybrid oocytes. **A**) Schematic shows HMGA2 degradation by Trim-Away after nuclear envelope breakdown. For simplicity, only a *musculus* oocyte is shown. **B**) Images show HMGA2-depleted *musculus*, hybrid, or *spretus* oocytes expressing TALE-mClover targeting major satellite and dCas9-mCherry with gRNA targeting minor satellite, fixed in meiosis I. Measurement brackets show representative lengths of pericentromeric satellites. Scale bars: 10 μm or 2 μm (inset); m: *musculus*, s: *spretus*. **C**) Quantification of pericentromeric satellite length in HMGA2-depleted *musculus*, hybrid, and *spretus* oocytes. Dotted line shows average length of major satellite in control hybrid oocytes reported in^18^. Each data point represents a single pericentromere; bars show mean and standard deviation; n=300-600 pericentromeres from 30 oocytes per condition from 3 independent experiments; Kruskal-Wallis test followed by Dunn’s multiple comparisons test, ****p<0.0001, ns: not significant. **D**) Images of control or HMGA2-depleted *musculus* oocytes in meiosis I, fixed, and stained with fluorescent anti-HMGA1 antibody. Scale bar: 10 μm. **E**) Quantification of HMGA1 intensity in control and HMGA2-depleted *musculus* oocytes. Each dot represents an average intensity of 10 chromosomes per single oocyte; bars show mean and standard deviation; n=23-29 oocytes from 3 independent experiments; two tailed-Mann-Whitney test.

### HMGA2 overexpression disrupts chromosome segregation in *spretus* but not hybrid oocytes

Given that HMGA2 abundance and major satellite copy number correlate (**Fig. 2**), and HMGA2 depletion disrupts major satellite packaging only in major satellite-rich *musculus* oocytes (**Fig. 3**), we wondered if protein abundance and satellite copy number must match to support meiotic chromosome segregation fidelity. We tested this by overexpressing HMGA2 in major satellite-poor *spretus* oocytes compared to hybrid oocytes with a much higher major satellite copy number. We first confirmed that HMGA2 mRNA microinjection increases HMGA2 abundance during meiosis I, with the expected pericentromere enrichment in *musculus* oocytes (**Fig. S3A and B**). We then increased HMGA2 abundance in *spretus* or hybrid oocytes, matured the oocytes *in vitro,* and counted the number of chromosomes in meiosis II-arrested eggs to assess the fidelity of chromosome segregation during the first meiotic division. Strikingly, HMGA2 overexpression increased aneuploidy in *spretus* but not hybrid oocytes (**Fig. 4A-D**). These findings show that high HMGA2 abundance interferes with meiotic chromosome segregation only when major satellite copy number is low.

**Fig. 4.**
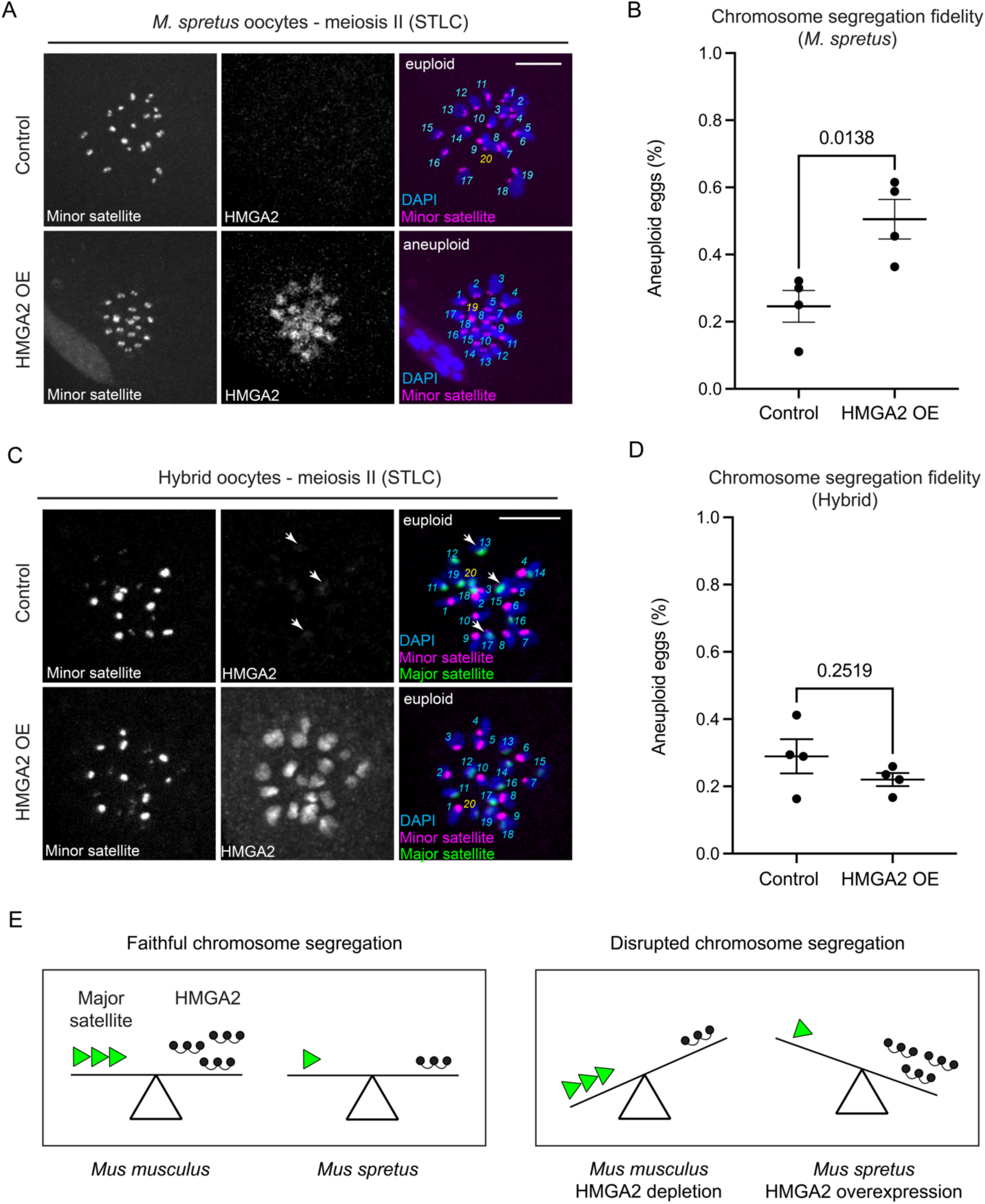
HMGA2 overexpression disrupts chromosome segregation in *spretus* but not hybrid oocytes. **A-D**) Images show *spretus* (A) or hybrid (C) oocytes expressing dCas9-mCherry with gRNA targeting minor satellite, without or without HMGA2 overexpression, fixed in meiosis II and stained for HMGA2. Hybrid oocytes also expressed TALE-mClover binding major satellite, with white arrows indicating increased localization of endogenous HMGA2 at major satellite. Numbers show chromosome counts; yellow numbers mark the last chromosome counted. Scale bars: 10 μm. Plots show the fraction of aneuploid eggs from *spretus* (C) or hybrid (D) oocytes. Each dot represents 9-28 (*spretus*) or 17-50 (hybrid) eggs per experiment, for 4 independent experiments. Bars show mean and standard error of the mean; unpaired two-tailed t-test. **E**) Model showing that chromosome segregation fidelity depends on balancing centromeric satellite copy number (e.g., major satellite) and the abundance of DNA shape-recognizing architectural proteins (e.g., HMGA2).

## Discussion

In this paper, we revealed functional and reciprocal constraints between the abundance of a highly conserved architectural protein and the copy number of a rapidly evolving pericentromeric satellite (**Fig. 4E**). We achieved this by first identifying HMGA2 as a major satellite-packaging protein whose abundance correlates with the copy number of that satellite in *musculus* vs *spretus* oocytes (**Figs. 1 and 2**). We tested functional constraints by experimentally manipulating both major satellite and HMGA2. Hybrid oocytes provide an intermediate satellite copy number between *musculus* (major satellite-rich) and *spretus* (major satellite-poor), while HMGA2 depletion or overexpression modulates protein abundance. HMGA2 depletion shows that HMGA1 is sufficient to package major satellite in hybrid oocytes, but both HMGA1 and HMGA2 are needed to package the higher major satellite copy number in *musculus* (**Fig. 3**).

Conversely, HMGA2 overexpression leads to chromosome segregation defects in major satellite-poor *spretus* but not hybrid oocytes (**Fig. 4**). Together, these results demonstrate that HMGA2 and major satellite abundance must match to support faithful chromosome segregation in female meiosis. HMGA2 depletion creates a mismatch when major satellite copy number is increased two-fold in *musculus* vs hybrid oocytes, while HMGA2 overexpression creates a mismatch with the small major satellite arrays in *spretus* vs hybrid oocytes.

Mechanistically, we propose that HMGA1 is sufficient to package major satellite at half the copy number found in *musculus*, explaining why HMGA2 depletion does not interfere with major satellite packaging in hybrid oocytes. Transient major satellite packaging defects were observed in *musculus*/*spretus* hybrid oocytes due to low levels of condensin II^17^, but we suspect that these defects are mechanistically distinct from the HMGA2-mediated packaging failure observed here in *musculus* oocytes. Unlike condensin II, HMGA2 recognizes a narrow DNA minor groove, crosslinks DNA molecules^55^, and forms nucleosome condensates^61^. These chromatin organizing properties can explain why HMGA2 overexpression increases aneuploidy in major satellite-poor *spretus* eggs. According to the heterochromatin sink hypothesis, heterochromatic DNA regions (e.g., pericentromeric satellites) modulate the availability of heterochromatin-binding proteins by sequestering them^62,63^. Similarly, the small major satellite arrays at *spretus* pericentromeres would not sequester the excess of HMGA2, which interferes with chromosome segregation fidelity. This effect is rescued in hybrid oocytes where excess HMGA2 is buffered by large major satellite arrays at *musculus* pericentromeres. Although the mechanism of HMGA2-mediated aneuploidy remains to be determined, an excess of HMGA2 might disrupt chromatin organization at minor satellite and interfere with kinetochore function. Altogether, our work expands the list of functions that DNA shape-recognizing architectural proteins play in genome integrity and organization^64–67^. Future work will focus on dissecting the role of architectural proteins during meiotic chromosome segregation.

Our findings support our model for the rapid divergence of centromeric satellites between closely related species (**Fig. 1A**). We propose that stochastic expansions of centromeric satellites from a common repeat library (e.g., via break-induced replication^35^) are tolerated only if they do not disrupt chromosome segregation. The strength of this selection is modulated by the abundance of DNA shape-recognizing architectural proteins that package the expanded satellites, allowing them to withstand spindle-pulling forces. The cost of satellite expansion is high in the absence of sufficient architectural proteins to package that satellite. Conversely, the cost is reduced if the packaging protein is abundant, and the expanded array persists. Under this model, HMGA2 abundance in *musculus* oocytes reduces the cost of major satellite expansion, and low HMGA2 abundance in *spretus* oocytes raises it, leading to differences in major satellite array sizes between *musculus* and *spretus*. A key next step in testing our satellite expansion model will be to manipulate the abundance of architectural proteins and monitor changes in centromeric satellite copy number over time. Finally, our findings suggest that centromeric satellites co-evolve with the abundance of highly conserved architectural proteins, complementing established models in which satellite divergence is coupled to amino acid changes in satellite-binding proteins^36,50,51,68–70^.

## Methods

### Mice

*Mus musculus* mice were purchased from Envigo/Inotiv (NSA, stock no. 033 corresponding to CF-1; used for oocyte experiments). *Mus spretus* mice were purchased from RIKEN BioResource Research Center (SPR2, RBRC00208). CF-1 females were crossed to *Mus spretus* males to generate hybrids. All mice used in this study were female at 2–9 months old. Mice used to test gene expression were age-matched. Mice were housed in controlled room-temperature conditions with minimal disturbances, a light–dark cycle of 12 h each, and humidity ranging from 30% to 70% depending on the season. All animal experiments were approved by the Institutional Animal Care and Use Committee of the University of Pennsylvania and were consistent with the National Institutes of Health guidelines (protocol no.: 804882).

### Oocyte collection and culture

Female mice were primed with pregnant mare somatic gonadotropin (*M. musculus*; 5 U per mouse; 367222; Calbiochem) or CARD HyperOva (*M. spretus* and *M. musculus* × *spretus* hybrid; 150 µl per mouse; Cosmo Bio USA; KYD-010-EX-X5) injected into the intraperitoneal cavity 38–48 h prior to oocyte collections to induce superovulation. The ovaries were isolated in M2 medium (M7167; Sigma-Aldrich Milipore) with 1.7 mM phosphodiesterase 3 inhibitor milrinone (M4659; 2.5 mM; Sigma-Aldrich Milipore) to maintain oocytes arrested in prophase I. The ovaries were dissected using a 27G size needle, and germinal vesicle oocytes were mechanically separated from cumulus cells using a mouth pipette with plastic 100-µm-diameter stripper tips (CooperSurgical; MXL3-100) and incubated in 5% CO2 (Airgas) for at least 1 h at 37.8 °C under mineral oil (9305; FUJIFILM Irvine Scientific) before microinjection.

### mRNA and plasmid preparation

HMGA2-OLLAS coding sequence with T7 promoter was synthesized as a gBlocks Gene Fragment (Integrated DNA Technologies). The pGEMHE-HMGB1-GFP and pGEMHE-HMGB2-GFP plasmids were generated by subcloning HMGB1 and HMGB2, respectively, from a cDNA library of an E12.5 (*Mus musculus x Mus spretus*) F1 hybrid embryo into a pGEMHE backbone with a C-terminal GFP tag. Inserts and backbones were linearized by PCR reaction using KAPA Polymerase HiFi HotStart ReadyMix (KK2602; Roche) and ligated using In-Fusion Snap Assembly (638948; Takara).

Primers were designed using SnapGene 8.0.3 (Dotmatics) software. Plasmids generated elsewhere: pIVT–dCas9–mCherry expressing catalytically dead Cas9, which targets the minor satellite through a gRNA that recognizes the 5′-ACACTGAAAAACACATTCGT-3′ sequence^71^; pTALYM3–TALE–mClover (kind gift from M.-E. Torres-Padilla; Addgene; no. 47874) expressing a TALE protein that recognizes the 5′-TGCCATATTCCACGT-3′ sequence of the major satellite repeat^72^. mRNA was prepared using an *in vitro* transcription kit: T7 mScript Standard mRNA Production System (C-MSC100625; CellScript) following the manufacturer’s instructions.

### Oocyte microinjection

Oocytes were microinjected with approximately 5 pl of mRNA in M2 medium with 1.7 mM milrinone and 3 mg ml^−1^ BSA at room temperature with a micromanipulator TransferMan 4r (Eppendorf) and picoinjector (Medical Systems Corp). Oocytes were then incubated in 30 μl drops of Chatot–Ziomek–Bavister (CZB) medium (MR019D; Thermo Fisher Scientific) under mineral oil (M5310; Sigma-Aldrich Milipore) at 37.8 °C and 5% CO2 (Airgas) for 3 h to allow protein expression. To deplete HMGA2, the Trim-away approach was used^73^. Briefly, milrinone-arrested germinal vesicle oocytes were microinjected with approximately 5 pl mix containing Trim21–OLLAS mRNA and 0.5 mg ml^−1^ of purified polyclonal normal rabbit IgG (used as control; 12-370; Sigma-Aldrich Millipore) or rabbit monoclonal anti-HMGA2 antibody (1:200; Cell Signaling 8179S, clone D1A7). RNA used: TALE–mClover 20-200 ng μl^−1^; Trim21–OLLAS 100-600 ng μl^−1^; dCas9–mCherry 15-75 ng μl^−1^; gRNA 2.7-15 ng μl^−1^; HMGA2–OLLAS 100 ng μl^−1^. Oocytes were then incubated for 3 h to enable mRNA expression before the cells were released by washing out milrinone. This was achieved by passing cells through five 100-μl drops of CZB medium. The polyclonal normal rabbit IgG antibody was passed through Amicon Ultra-0.5 100-K columns (Sigma-Aldrich Milipore, UFC5100625) to remove preservatives and glycerol following an established protocol^74^. IGEPAL at 0.05% final concentration was added to facilitate injections of viscous mRNA and protein solutions.

### Oocyte immunofluorescence

Oocytes were fixed in 2% paraformaldehyde and 0.1% Triton X-100 dissolved in PBS (pH 7.4) for 20 min at room temperature, then placed in 0.1% Triton X-100 for 15 min at room temperature, placed in blocking solution (PBS with 0.3% BSA and 0.01% Tween-20) overnight at 4 °C, incubated for 1 h with primary antibody in blocking solution, washed three times for 15 min each, incubated for 1 h with secondary antibody, washed three times for 15 min each and mounted in Vectashield with DAPI (H-1200; Vector) to visualize chromosomes. Rabbit polyclonal anti-HMGA2 antibody (1:200; ab97276; Abcam) was used to detect HMGA2. Rat monoclonal anti-OLLAS (1:200; clone L2; NBP1-06713; Novus Biologicals) antibody was used to detect HMGA2-OLLAS. AlexaFluor 488-conjugated rabbit monoclonal anti-HMGA1 antibody (1:500; EPR7839 clone; ab204667; Abcam) was used to detect HMGA1 in HMGA2-depleted oocytes.

AlexaFluor 488-conjugated rabbit monoclonal anti-HMGA2 antibody (1:100; EPR18114; Abcam; ab317357) was used to detect HMGA2 in HMGA2-depleted oocytes. The secondary antibodies used: anti-rabbit AlexaFluor 488 (1:500, Invitrogen), anti-rat AlexaFluor 488 (1:500, Invitrogen), anti-rabbit AlexaFluor 647 (1:500, Invitrogen), and anti-rat AlexaFluor 647 (1:500, Invitrogen).

### Reverse transcription-quantitative polymerase chain reaction (RT-qPCR)

Total RNA was extracted from 30 age-matched *Mus musculus*, hybrid, or *Mus spretus* oocytes using the Arcturus PicoPure RNA Isolation Kit (KIT0204; Applied Biosystems). Extracted RNA was treated with the RNase-Free DNase Set (79254; Qiagen) and reverse transcribed using the SuperScript IV First-Strand cDNA Synthesis System (18091050; Invitrogen). Primers for the control genes *Hprt1* and *Sptbn1* were selected based on previous studies^75,76^, verified to recognize transcripts from both *musculus* and *spretus* using the Ensembl database. Primers to detect HMGA1 and HMGA2 transcripts were designed using PrimerQuest (IDT). HMGA1 primers recognize isoforms *Hmga1a* and *Hmga1b* in both *musculus* and *spretus*. Primers for HMGA2 also recognize *musculus* and *spretus Hmga2* transcripts, which both encode: MSARGEGAGQPSTSAQGQPAAPVPQKRGRGRPRKQQQEPTCEPSPKRPRGRPKGSKNKSPSKAAQKKAETIGEKRPRGRPRKWPQQVVQKKPAQETEETSSQESAEED.

Primer sequences used: *Hmga1*: 5’-GAAGTGCCAACTCCGAAGA-3’, 5’-CTCTTCCTCCTTCTCCAGTTTC-3’; *Hmga2*: 5’-AGCAAGAGCCAACCTGTG-3’, 5’-CTTTCTTCTGGGCTGCTTTAGA-3’; *Hprt1*: 5’-GCTTGCTGGTGAAAAGGACC TCTCGAAG-3’, 5’-CCCTGAAGTACTCATTATAGTCAAGGGCAT-3’; *Sptbn1*: 5’-TCTAATGGTTACTTGCTTGTGCG-3’, 5’-CAATAGTTACAGTGACAGAGAATGC-3’.

Importantly, the primers do not recognize the transcript for the truncated version of HMGA2, a likely pseudogene^77^. Each qPCR experiment was conducted using a QuantStudio 3 Real-Time PCR System (A28567; Applied Biosystems) and a MicroAmp Optical 96-Well Reaction Plate with Barcode (4306737; Applied Biosystems). qPCR conditions were optimized according to the manufacturer’s instructions and by testing different melting temperatures and template concentrations. For each reaction, 2μL of template DNA (or water for no template controls) was added to a mix of 10μL of SYBR Green Universal Master Mix (4309155; Applied Biosystems), 1μL of 5μM forward primer, 1μL of 5μM reverse primer, and 6μL water. Data were analyzed using the QuantStudio Design and Analysis Software v2, Relative Quantification Analysis Module (ThermoFisher). Three biological and three technical repetitions were done for each gene and species. The mean value of these triplicates was used to calculate Cq values.

### Image acquisition

Images of oocytes were collected as *z* stacks (31 or 41 slices) at 0.5-μm intervals to visualize the entire meiotic spindle using a confocal microscope (DMI4000B; Leica) equipped with a 63×, 1.3 NA glycerol-immersion objective lens, an *xy*-piezo *z* stage (Applied Scientific Instrumentation), a spinning disk (CSU10; Yokogawa) and an electron multiplier charge-coupled device camera (ImageEM C9100-13; Hamamatsu Photonics), controlled by MetaMorph 7.5 software (Molecular Devices). Excitation was carried out with a Vortran Stradus VersaLase 4 laser module with 405-, 488-, 561- and 639-nm lasers (Vortran Laser Technology). Panels of microscopy images were prepared using ImageJ 1.54f.

### Image analysis

To quantify HMGA2, HMGB1, and HMGB2 enrichment at pericentromeres in *M. musculus*, hybrid, and *M. spretus* bivalents, average intensity projections of the z-slices for each bivalent were used. Fifteen-pixel-long and one-pixel-wide lines were drawn to measure intensities at major (*M. musculus*) or minor (*M. spretus*) satellites, chromosome arm, and the nearby cytoplasm. Lines were centered on the satellite region, and the intensities of the middle pixels within each region were compared (3^rd^ pixel – cytoplasm, 8^th^ pixel – satellite, and 13^th^ pixel – chromosome arm). Values were obtained for ten (HMGA2) or five (HMGB1 and HMGB2) randomly selected, non-overlapping bivalents for each cell. HMGA2 depletion efficiency was calculated based on maximum projection images. A DAPI-based mask was created for each image using the Li threshold algorithm for control or HMGA2-depleted cells and ImageJ 1.54f software plug-in Analyze Particles (size 3-infinity; circularity 0.00–3.00). Any image artifacts were manually removed from the mask. Mean HMGA2 intensity within the mask was measured, and a 50 × 50-pixel box drawn at the cytoplasm adjacent to chromosomes was used to subtract the cytoplasmic background. To quantify the impact of HMGA2 depletion on satellite packaging for *M. musculus*, hybrid, and *M. spretus* bivalents, maximal intensity projections for the entire range of the z-stack were used.

Lines were drawn from the beginning of the satellite, as indicated by the border between DAPI and satellite signals, to the end of the satellite, as indicated by the furthest point on the border between the cytoplasm and satellite signal. Values were obtained for ten randomly selected, non-overlapping bivalents for each HMGA2-depleted cell. To quantify HMGA1 intensity in HMGA2-depleted cells, maximum intensity projections for the entire range of the z-stack were used. Circular six-pixel diameter regions were drawn to measure intensities at the chromosome arm for each bivalent, and a 50 x 50-pixel box was drawn on the nearby cytoplasm to measure background values for each cell. Values were obtained for ten randomly selected, non-overlapping bivalents for each cell. To quantify the intensity of HMGA2 at pericentromeres in *M. musculus*, hybrid, and *M. spretus* bivalents, average intensity projections of the z-slices for each bivalent were used. Circular six-pixel-diameter regions were drawn to measure intensities at major and minor satellites and at non-pericentromeric chromatin. Values were obtained for ten randomly selected, non-overlapping bivalents for each cell stained for HMGA2. To quantify aneuploidy of hybrid and *spretus* oocytes, the number of chromosomes in meiosis II eggs was counted using the DAPI, major satellite (hybrid only), and minor satellite signals. Both maximal intensity projections and individual z-slices were used.

### AlphaFold 3 analysis

Protein-DNA binding modeling was done using AlphaFold 3 (AlphaFold Server; https://alphafoldserver.com)^78^. An *in silico* competition assay between major and minor satellites for protein binding was designed by creating a chimeric DNA molecule. First, to mimic the previously established differences in DNA shape between major and minor satellites^18^, HMGA1 protein binding was modeled using the full consensus sequence of a major (234 bp) or minor (120 bp) satellite repeat^79,80^. A contiguous sequence that included all sites where HMGA1 AT-hook motifs were bound was defined as the seed sequence. These seeds were extended so that the major satellite sequences had 5 AT-stretches across 45 bp, while those for the minor satellite had 3, representing the proportional density of AT-stretches for each major and minor satellite repeat^18^. Second, a 30 bp guanine linker was added to separate the major and minor satellite fragments.

Third, protein binding to the created chimeric DNA molecule was modeled. All five output models were inspected using Pymol (The Pymol Molecular Graphics System, Version 3.1.3.1, Schrodinger, LLC) to determine binding preference. The assay was repeated with the relative positions of major and minor satellite fragments swapped to ensure the position of each binding site within the chimeric DNA molecule did not affect the modeling. Only cases in which all ten models for both chimeric sequences agreed were considered as evidence of preference. The same chimeric DNA molecule was used to model the binding of HMGA1, HMGA2, HMGB1, and HMGB2.

### Statistical analysis

Experiments were done in three biological replicates unless specified. Statistical analyses were performed using Prism 11 (GraphPad). Sample size choice was guided by the number of cells and bivalents that captured the variation between samples and across conditions. Except for the experiment testing the effects of HMGA2 overexpression on aneuploidy (Fig. 4), the investigators were not blinded during data collection and quantification because the phenotypic changes were readily distinguishable across the analyzed conditions. For all experiments, the cells or bivalents analyzed were selected at random.

## Resource availability

### Lead contact

### Materials availability

All unique reagents in this study are available upon request to the lead contact.

### Data and code availability

- All data reported in this paper, which includes raw image files, will be shared by the lead contact upon request.
- This paper does not report any original code.
- Any additional information required to reanalyze the data reported in this paper is available from the lead contact upon request.

## Acknowledgements

We thank all the members of the M. Lampson and M. Levine labs at the University of Pennsylvania for comments on the data, and members of the Philadelphia Chromosome Club for discussions on our findings. Special thanks to Jacob Cote and Amber Ridgway for help with RT-qPCR experiment design and setup. Funding was from NIH grant GM122475 (M.A.L.).

## Author Contributions

D.D. and M.A.L. conceived the project. D.D. and K.B. generated reagents and performed experiments. All authors analyzed the data. D.D. wrote the paper and prepared the figures. K.B. wrote the Methods section. D.D. and M.A.L. edited the paper. D.D. and M.A.L. supervised the project.

---

**Supplementary Fig. 1.**
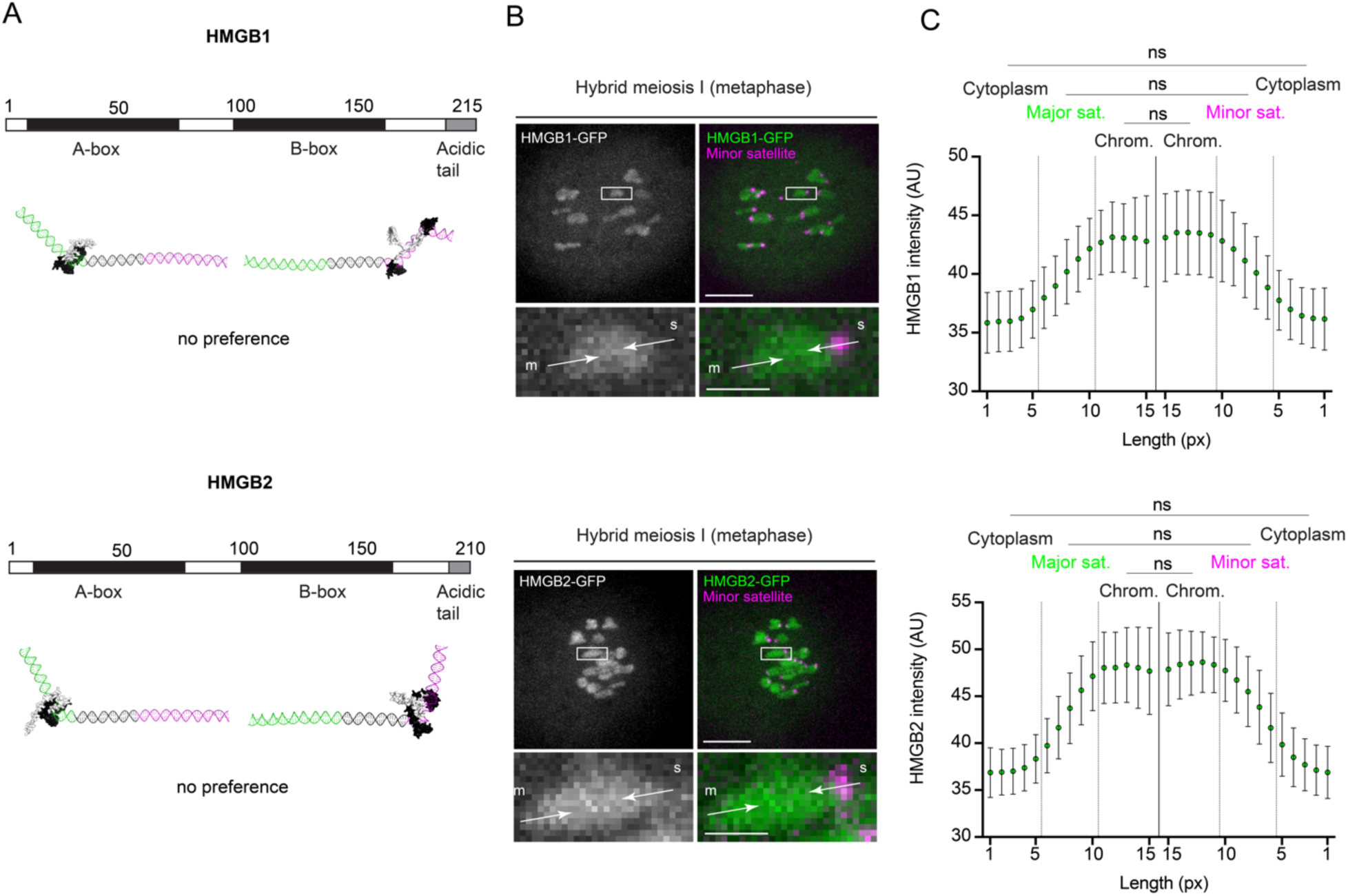
HMGB1 and HMGB2 do not show a preference for major satellite. **A**) AlphaFold3 modeling of HMGB1 (top) and HMGB2 (bottom) binding to a model DNA molecule containing major (green) and minor (magenta) satellite fragments separated by a poly-guanine linker (black). **B**) Images show hybrid oocytes in meiosis I expressing dCas9-mCherry with gRNA targeting minor satellite, together with either GFP-tagged HMGB1 (top) or HMGB2 (bottom). Arrows show line scans along homologous chromosomes. Scale bars: 10 μm or 2 μm (inset); m: *musculus*, s: *spretus*. **C**) Quantification of HMGA1 (top) or HMGB2 (bottom) intensity in hybrid oocytes, averaged over multiple line scans. Bars show standard deviation; n=50 line scans from 5 oocytes; Kruskal-Wallis test followed by Dunn’s multiple comparisons test, ns: not significant.

**Supplementary Fig. 2.**
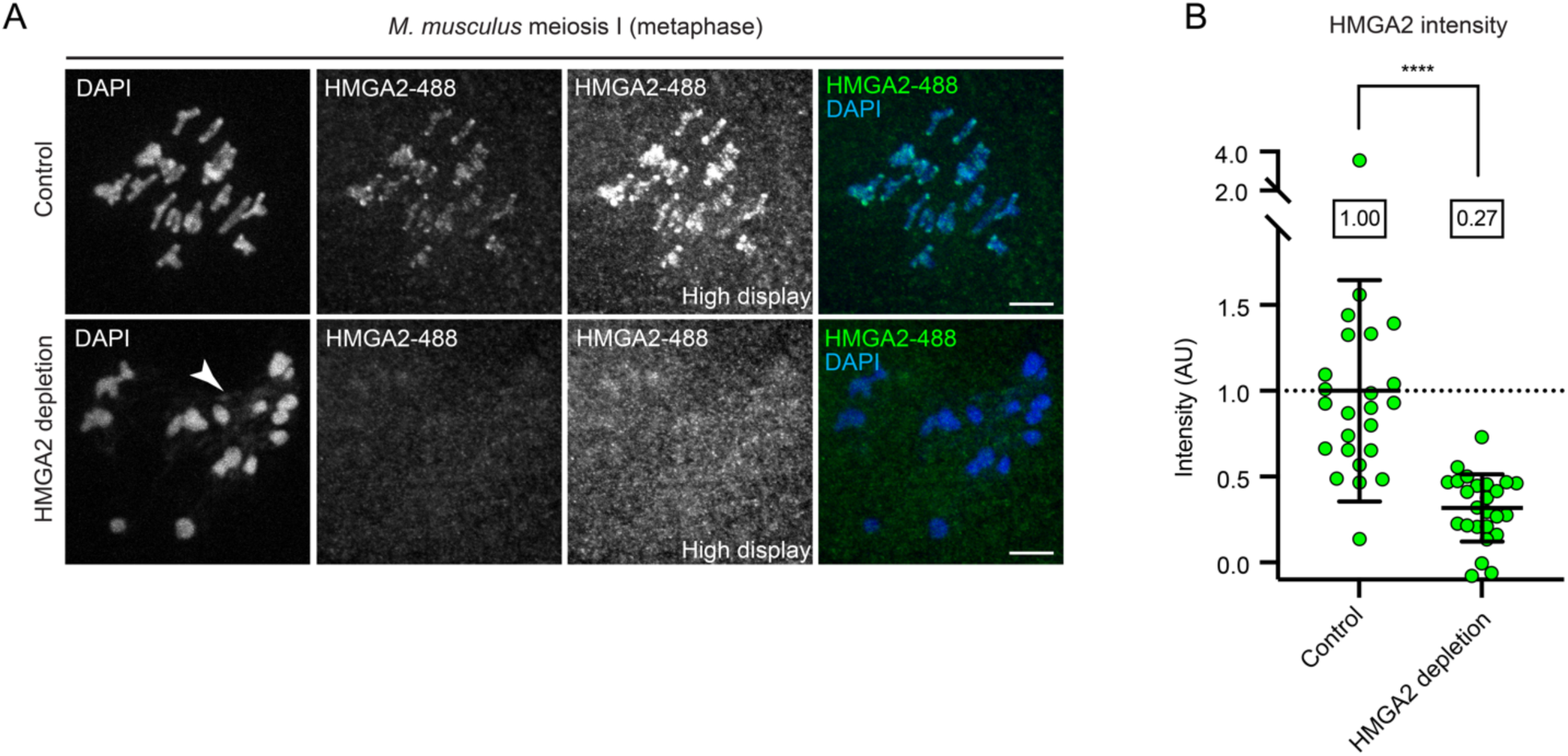
Trim-away efficiently depletes HMGA2 protein. **A**) Images show control or HMGA2-depleted *musculus* oocytes, fixed in meiosis I and stained with fluorescent anti-HMGA2 antibody. **B**) Quantification of HMGA2 intensity in control and HMGA2-depleted *musculus* oocytes. Each dot represents the average intensity per single oocyte. Bars show mean and standard deviations; n=24-25 oocytes from 3 independent experiments; two-tailed Mann-Whitney test, ****p<0.0001.

**Supplementary Fig. 3.**
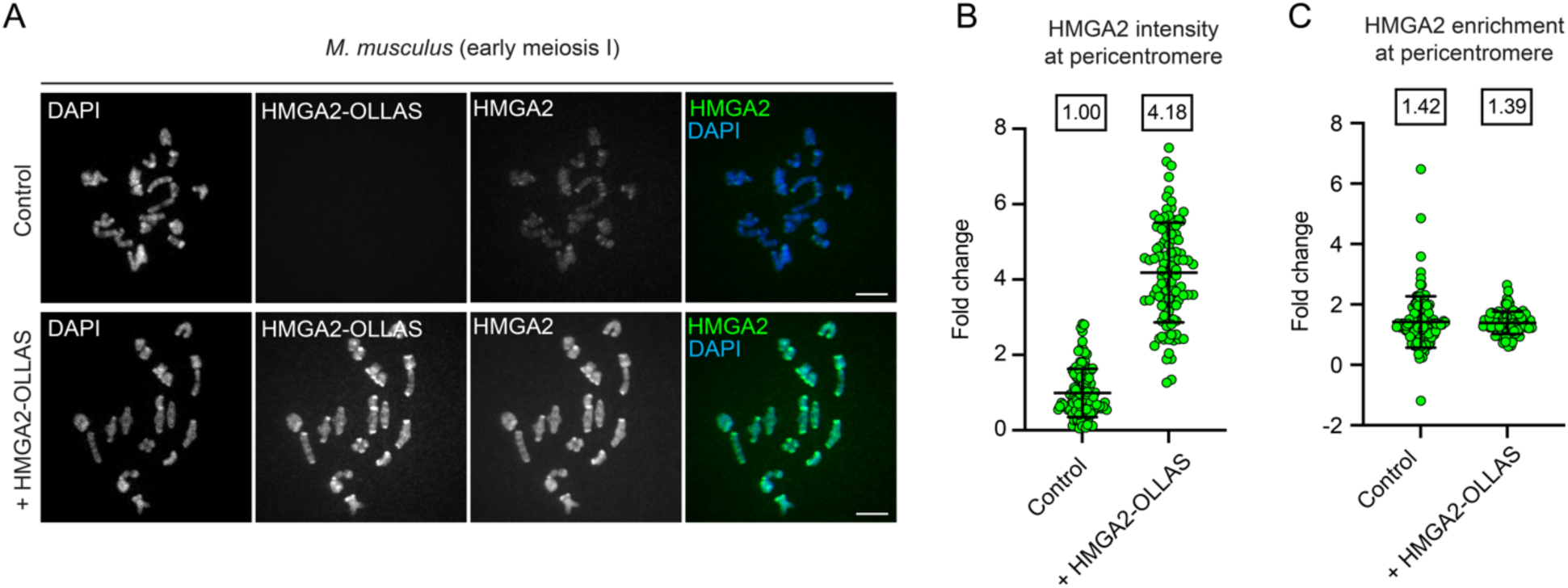
HMGA2-OLLAS microinjection increases HMGA2 levels at pericentromeres. **A**) Images show *musculus* oocytes with or without expression of HMGA2-OLLAS, fixed in meiosis I and stained for both HMGA2 (labeling both endogenous and exogenous protein) and the OLLAS epitope tag (exogenous protein). **B** and **C**) Quantification of total HMGA2 intensity in control and HMGA2-OLLAS-expressing *musculus* oocytes (B) or enrichment of HMGA2 at pericentromeres compared to chromosome arms (C). Each dot represents a single pericentromere; bars show mean and standard deviation; n=100-120 from 10-12 oocytes.

